# Establishment of porcine ovarian follicles treated with t-BHP for an in vitro aging model

**DOI:** 10.1101/2022.09.23.509190

**Authors:** Peihua Shi, Jinchun Gao, Shunran Zhao, Wei Xia, Junjie Li, Chenyu Tao

**Affiliations:** College of Animal Science and Technology, Hebei Agricultural University, Baoding, Hebei Province, China

## Abstract

Ovarian aging is closely associated with low female fertility. Excessive oxidative stress can induce ovarian senescence and follicular atresia, thereby reducing reproductive performance. In this study, intact antral follicles of pigs were treated with tert-butyl hydroperoxide (t-BHP) to simulate aging stimulation conditions. The results showed that 200 μM t-BHP induced the senescence-related phenotype of antral follicles, which was time-dependent and changed significantly after 6 h of treatment. t-BHP can induce an increase in reactive oxygen species (ROS) and senescence mediated by Caspase-3, P53, Foxo1, and SOD. Senescence-associated β-galactosidase (SA-β-Gal) staining revealed an increase in the number of positive cells. Enzyme-linked immunosorbent assays (ELISA) showed that the level of reactive oxygen species increased, and that the ratio of progesterone (P_4_) to estradiol (E_2_) increased. Finally, the findings of this study showed that the relative expression levels of Caspase-3, P53, Foxo1 mRNA, and protein increased, while the relative expression levels of SOD decreased. In conclusion, treatment with 200 μM of t-BHP effectively induced follicular senescence at 6 h. Therefore, the in vitro treatment of follicles with 200 μM t-BHP for 6 h may be a feasible in vitro culture model to simulate ovarian senescence in gilts.

## Introduction

Epigenetics influences a variety of existing functions in organisms, including development, differentiation, stress, aging, and some pathological states. Epigenetics can alter gene expression in many ways, including DNA methylation, histone modification, genomic imprinting, chromosome inactivation, and non-coding RNA. Aging is regularly accompanied by an imbalance of homeostasis and decreased autophagy clearance of cells, mainly due to the dysfunction of protein and mitochondria which induces continual inflammation, and eventually, anxious inflammation[1, 2]. Cellular senescence is a state induced by stress and certain physiological processes, and is characterized by secretion, DNA damage, oxidative stress, prolonged cell cycle of metabolic changes, and usually irreversible cell cycle termination[3]. Cells have a restricted ability to replicate. Once a threshold is reached, the cells are no longer able to divide, even with increased stimulation. Once a wide variety of proliferating cells in a tissue undergo phone senescence, the regeneration ability of the tissue is reduced. Cellular senescence is regularly related to age cause a lack of regenerative ability is a marker of tissue senescence[4]. The number of senescent cells increases with age in multiple tissues[5-7].

In female mammals, aging has an extensive impact on reproductive performance. Ovarian senescence is a process in which the potential of the ovaries to produce fully functional gametes is progressively weakened. Ovarian senescence includes the non-stop loss of follicles and a limit in the whole quantity of follicles, which consequently leads to a gradual decrease in female fertility. In addition, due to a decrease in ovarian follicle cells, ovarian aging leads to the failure of the ovary to produce adequate sex hormones to maintain normal physiological features in animals or humans[8, 9]. Therefore, the continuous decline of follicle quantity with increasing age eventually leads to hormone imbalance and an irregular ovarian cycle. Additionally, aging ovaries also produce poor-quality oocytes, which can lead to developmental stagnation, aneuploidy, incapability to implant embryos, and abortion[10-12]. Granulosa cells and corpus luteum cells of follicles are necessary sources of estrogen and progesterone for female physiological activities. A permanent increase in the arrest of granulosa cells might also inhibit the capability of ovarian follicles to mature, mainly causing atresia of immature follicles.

The reproductive physiology of sows significantly changes after aging. On one hand, fecundity decreases, and the elimination rate increases. However, the decrease in dominant sows will result in serious losses in production efficiency. The ovaries of pigs typically develop unexpectedly from 72–165 days of age. HE staining for atresia regulation of follicles in the ovarian tissues of pigs confirmed that in the course of oocyte proliferation, spare oocytes were apoptotic. In the follicular stage, atresia of a wide range of primordial follicles and dominant follicles were observed, and the total primordial follicle population confirmed constant atresia[13]. Atresia of follicles at all stages can be observed in the fast follicular boom phase, with the degeneration of primordial follicles being the most significant. Some studies have shown that TUNEL staining can be used to examine the atresia of a large variety of primordial follicles and principal follicles in the oocyte proliferation stage, and that atresia is derived from the apoptosis of follicular oocytes[14]. Atresia can occur in all stages of follicle development. Atresia of the main follicles is typically prompted by the apoptosis of oocytes, and some are accompanied by the simultaneous apoptosis of oocytes and follicular cells. Atresia of secondary follicles was once broadly thought to be prompted by the apoptosis of granulosa cells. With the rapid increase in the length of follicles and the rapid boom of follicles, atresia of follicles at all ranges becomes more obvious.

With the enhanced survival of patients after treatment with chemotherapy for malignant tumors and immune diseases, it has been reported that chemotherapeutic drugs such as cyclophosphamide, cisplatin, tripterygium wilford, and hydrocortisone affect ovarian function and precipitate reproductive issues. There is an increased frequency in the use of chemical strategies to manage diseases. Cyclophosphamide, one of common chemotherapeutic drugs, has an exceptionally poisonous impact on the ovary. It works by inhibiting the synthesis of DNA, RNA, and protein, inflicting gonadal injury[15]. Because of the limited technology of ovarian culture in vitro and most of the chemical methods used to establish mouse models[16], aging models of porcine ovaries are rare[17, 18]. Most in vitro aging models are constructed using cells, and oxidative damage is one of the most commonly used methods to construct aging models. Oxidative stress-inducing conditions were adjusted for experimental purposes using different cells. Human skeletal myoblasts were treated with 1 mmol/L hydrogen peroxide for 30 min to investigate the regulatory effect of the tocotrienol-rich fraction on senescence[19]. Another study reported that t-BHP could induce the aging of mouse hematopoietic stem cells, and the percentage of aging cells reached 57.92% ± 4.24% after 6 h[20]. It is urgent to establish an in vitro aging model to study the biological function of ovarian aging because it is difficult to obtain in high-parity sow samples. Therefore, in this study, an aging model of follicles in vitro was established, which provided experimental data for the study of the mechanism of delaying aging on reproductive physiology.

## Materials and Methods

The basal medium DMEM/Ham’s F-12 (DMEM/F12) was purchased from Thermo Fisher Scientific. PBS was purchased from HyClone Laboratories Inc. (Logan, UT, USA). TRIzol was purchased from TIANGEN (Beijing,China). The reverse transcription kit was purchased from Ta’KaRa (Dalian,China) and SYBR Green was purchased from Servicebio Biotechnology (Wuhan, China). The cell senescence β-galactosidase staining package was purchased from Beyotime Biotechnology (C0602, Shanghai, China). The porcine estradiol enzyme-linked immunosorbent assay package was purchased from Yuanmu Biotechnology (YM-S2619, Shanghai, China). The porcine progesterone enzyme-linked immunosorbent assay package was purchased from Yuanmu Biotechnology (YM-S3357, Shanghai, China). The porcine ROS ELISA package was purchased from Yuanmu Biotechnoogy(YX-181519P, Shanghai, China).

### Ovaries source

The ovaries of sexually mature pigs had been amassed from a neighborhood slaughterhouse in Baoding, Hebei Province, and placed in ordinary saline supplemented with 100 U/mL penicillin and 100 g/mL streptomycin at 37 °C, stored in vacuum flasks, and introduced again to the laboratory as quickly as possible.

### Screening and culture of intact antral follicles

The ovaries were sprayed with 75% ethanol to create a sterile environment and then rinsed two to three times with 37 °C saline containing antibiotics until water was clear. The ovaries were placed in a preheated bottle containing saline and antibiotics. The isolated follicles without red/corpus luteum, with uniform and transparent texture and dense distribution of antral follicles, were selected, and the diameter of the isolated follicles was measured using a ruler. The intact follicles measuring 3–6 mm in diameter, with the follicular cavity, full follicular cavity, rich blood vessels, and clear follicular fluid without dark spots were selected and seeded in 24-well plates; one follicle per well, at 37 °C, 5% CO2, and incubated in a constant humidity incubator. Morphological changes in the follicles were observed and recorded, such as whether the color of the follicles turned gray, the wall of the blood vessels decreased, the follicular fluid showed flocculant deposits, and dark spots were present on the inner wall of the follicles.

### Treatments

The cells were divided into three groups according to treatment: control, time processing, and t-BHP-treated groups. Pre-treatment for 24 and 36 h was used to evaluate the appropriate culture conditions for maintaining healthy follicles in the current study. Follicles in the t-BHP-treated group were treated with 200 μM t-BHP for 1, 2, 6, and 12 h to simulate the oxidative stress-induced senescence microenvironment.

### ELISA detection of hormone in follicle fluid

A total of 50 μL pattern was placed in every well and incubated at 37 °C for 30 min. Each properly was once brought with a 50 μL enzyme-labeled reagent and incubated at 37 °C for 30 min. A50 μL shade developer was added to every well, along with B50 μL shade developer, and the solution was mixed with mild shaking and allowed to react in the dark at 37°C for 15 min. Then, 50 μL quit answer was brought to every properly to the reaction. The OD density of each well was measured at 450 nm.

### Detection of ROS level in follicular fluid by ELISA

Then, 50 μL pattern and 100 μL of detection antibody were added into every well and incubated at 37 °C for 60 min. Substrates A and B (50 μL each) were added into every well and the solutions were incubated in the dark at 37 °C for 15 min. 50 μL stop buffer were introduced into every well. The OD was measured at a wavelength of 450 nm within 15 min.

### Ovarian granulosa cell acquisition

Granulosa cells were collected after the follicle culture was completed. The follicles were removed from the 24-well plate with sterile ophthalmic tweezers; the follicle membrane was cut with a sterile surgical blade and follicle fluid was collected. The follicle fluid was filtered through a cell sieve and inoculated into a 6-well plate, and cell growth was observed by adding 15% DMEM/F 12 complete medium.

### SA-β-Gal cytochemical staining

Cells of the control group and treatment groups were collected, washed twice with phosphate-buffered saline (PBS), and fixed with fixing solution at room temperature for 15 min. Then, the cells were washed twice with PBS and incubated overnight with freshly prepared staining solution at 37 °C without CO_2_ for 12 h, as recommended by the manufacturer. The percentage of SA-β-Gal cells was determined by counting the number of blue cells under bright field illumination and total number of cells in the same field under phase contrast.

### ROS staining

Briefly, DCFH-DA probes were directly added to serum-free medium to be diluted to the working concentration (final concentration: 10 μM), and 1 mL of diluted DCFH-DA solution was added after the medium in the six-well plate was discarded. The medium was incubated for 30 min at 37°C, discarded, and washed with PBS twice before observation under a laser confocal microscope.

### RNA extraction and Quantitative real-time PCR (qRT-PCR) analysis

The total RNA of the cells was extracted using TRIZOL. RNA concentration and quality were determined using a spectrophotometer NanoDrop 2000 (Thermo Fisher Scientific, US). Samples with RNA concentrations between 500 and 1,000 ng/mL and OD 260/280 value between 1.8 and 2.0 were selected for further analysis. According to the reverse transcription kit instructions, 1 mg of RNA was transformed into cDNA by reverse transcription. qRT-PCR was performed using the SYBR Green method. The differential abundance of the mRNA of genes relative to GAPDH was determined using the 2^-ΔΔCT^ method. Primers were synthesized using GENEWIZ (Jiangsu, China) (Table 1).

**Table 1.**
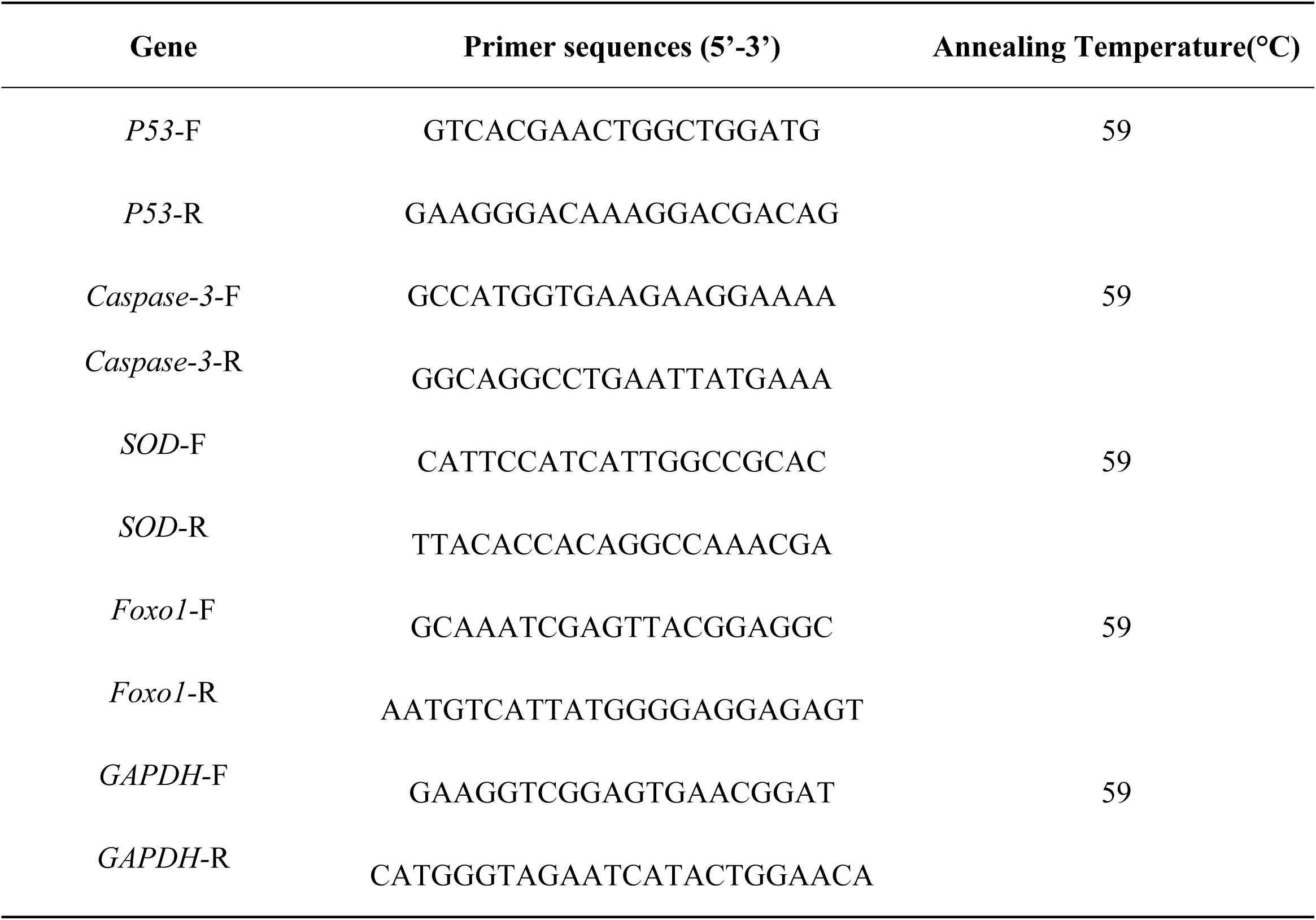
Primers used in this study.

### Western blot analysis

Western blot analysis was performed as described previously[21]. The membranes were visualized using electrochemiluminescence. The data were analyzed with Quantity One. The antibodies used are shown in Table 2.

**Table 2.** Antibodies used in this study.

| Antibody | Host Species | Dilution | Molecular weight (KD) | Vendor |
| --- | --- | --- | --- | --- |
| GAPDH | Mouse | 1: 2000 | 37 | Servicebio |
| Caspase-3 | Rabbit | 1: 1000 | 35 | Servicebio |
| Foxo1 | Rabbit | 1: 1000 | 78 | Servicebio |
| P53 | Rabbit | 1: 1000 | 55 | Servicebio |
| SOD2 | Rabbit | 1: 1000 | 25 | Servicebio |
| Anti-rabbit | Goat | 1 : 5000 | - | Servicebio |
| Anti-mouse | Goat | 1 : 5000 | - | Servicebio |

### Statistical analysis

Data are shown as the mean ± standard error of the mean (SEM) from three independent experiments. The experimental data were sorted using Excel 2007 software, and the diagram was drawn using Prism 9. Statistical significance was determined using the SPSS software version 26.0 (SPSS V26.0). One-way analysis of variance (ANOVA) was performed, and pairwise comparisons were performed using the LSD method. Statistical significance was set at p < 0.05.

## Results

### Quality identification of porcine antral follicles

To verify the accuracy of judging the morphology of porcine isolated follicles, healthy follicles were selected according to size (3–5 mm in diameter) (Fig 1A), and the follicles of the three groups were cultured and observed under a microscope.

**Fig 1.**
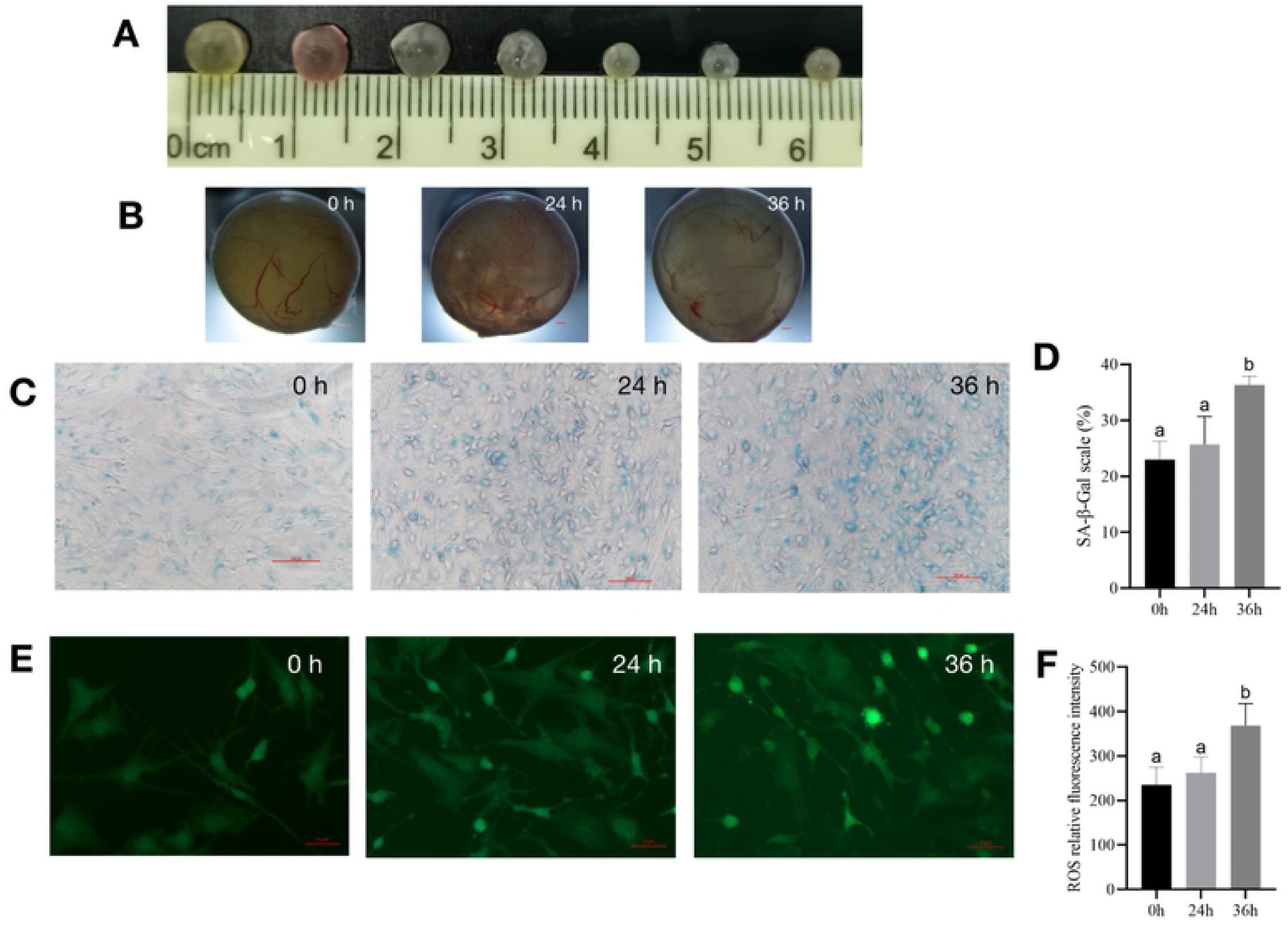
The culturing system of porcine antral follicles is feasible within 24 h. (A) The antral follicles with a diameter of 3–5 mm was selected. (B) The antral follicles were cultured in vitro for 24 and 36 h, and the bar represents 10 μm. (C) SA-β-Gal staining of the granulosa cells. The bar represents 20 μm. (D) Statistics of positive cells at different time points. P<0.05. (E) ROS fluorescence intensity staining of follicles in each group. The bar represents 20 μm. (F) Relative ROS levels. P<0.05.

The morphological characteristics of the follicles at each stage were as follows: the follicle wall of the control group was intact, even, and compact, with a little pink or a few yellow regions, and the capillaries were well distributed and bright red. Follicular fluid was clear, and there were no shed granulosa cells in the follicular cavity. In the 24-h group, the outer layer of the follicular wall was intact, the basement membrane was partly degraded, the capillary decreased to different degrees, the follicular fluid was clear, and the follicular cavity was clean and without lumps. In the 36-h group, the outer layer of the follicle wall was intact with fewer capillaries, a few granulosa cells exfoliated into the follicular cavity, and a small of flocculent deposits appeared in the cytoplasm (Fig 1B). Granulosa cells in the parietal layer of the follicles were stained with SA-β-Gal to observe cellular senescence. It was found that the cells in the control group were short and round with close intercellular connections, while the staining of positive cells was deepened and the intercellular connection was not closed; the cells were partially deformed (Fig 1C), and the ratio of positive cells was found to not be significantly higher at 24 h than in the control group, and significantly higher at 36 h (Fig 1D).

The results fluorescence staining showed that ROS in the 24 h group was not significantly higher than that in the control group, but significantly increased after 36 h (Fig 1E, F).

### Changes in progesterone and estrogen in follicles

The results of hormone level measurement showed that the concentration of E_2_ decreased significantly and the concentration of P_4_ increased significantly after 36 h, but there was no significant difference between the two groups at 24 h. The P_4_/E_2_ ratio increased significantly with an increase in follicular culture time (Table 3). This shows that atresia did not occur in the follicles during in vitro culture. Based on the above results, it can be confirmed that 24 h of in vitro culture of follicles can be used in follow-up experiments.

**Table 3.** The changes of hormone level in follicles.

| Processing time | Progesterone concentration<br>(pmol/L) | Estradiol<br>concentration<br>(pmol/L) | P<sub>4</sub>/E<sub>2</sub> |
| --- | --- | --- | --- |
| 0 h | 140.00±19.51<sup>a</sup> | 218.70±4.94<sup>a</sup> | 0.64±0.08<sup>a</sup> |
| 24 h | 190.00±3.21<sup>b</sup> | 198.52±7.04<sup>b</sup> | 0.96±0.02<sup>b</sup> |
| 36 h | 223.33±14.70<sup>b</sup> | 149.58±4.26<sup>c</sup> | 1.50±0.13<sup>c</sup> |

Different letters indicate statistically significant differences (P < 0.05).

### The optimal time of follicular senescence induced by t-BHP

To verify whether t-BHP treatment at 200 μM for 1 h, 2 h, 6 h and 12 h induced follicular senescence, we did further experiments with the following treatments: a control group without t-BHP and, and treatment with 200 μM sodium butyl peroxide for 1 h, 2 h, 6 h and 12 h, respectively. After treatment, the follicles were cultured in drug-free medium for 24 h, and granulosa cells were collected and cultured until they adhered to the wall.

The follicular phenotype showed intact follicular walls, visible blood vessels, and a clear follicular cavity 1 and 2 h after culture (Fig 2A). In the 6-h group, follicular blood vessels decreased, and a small number of dark spots appeared in the follicular cavity (Fig 2A). However, at 12 h, the blood vessels in the wall of the follicle disappeared, and the color of the follicle gradually became gray, with flocculent deposits in the lumen. The granulosa cells in the parietal layer broke off into clumps, which were significantly different from those in the control group (Fig 2A).

**Fig 2.**
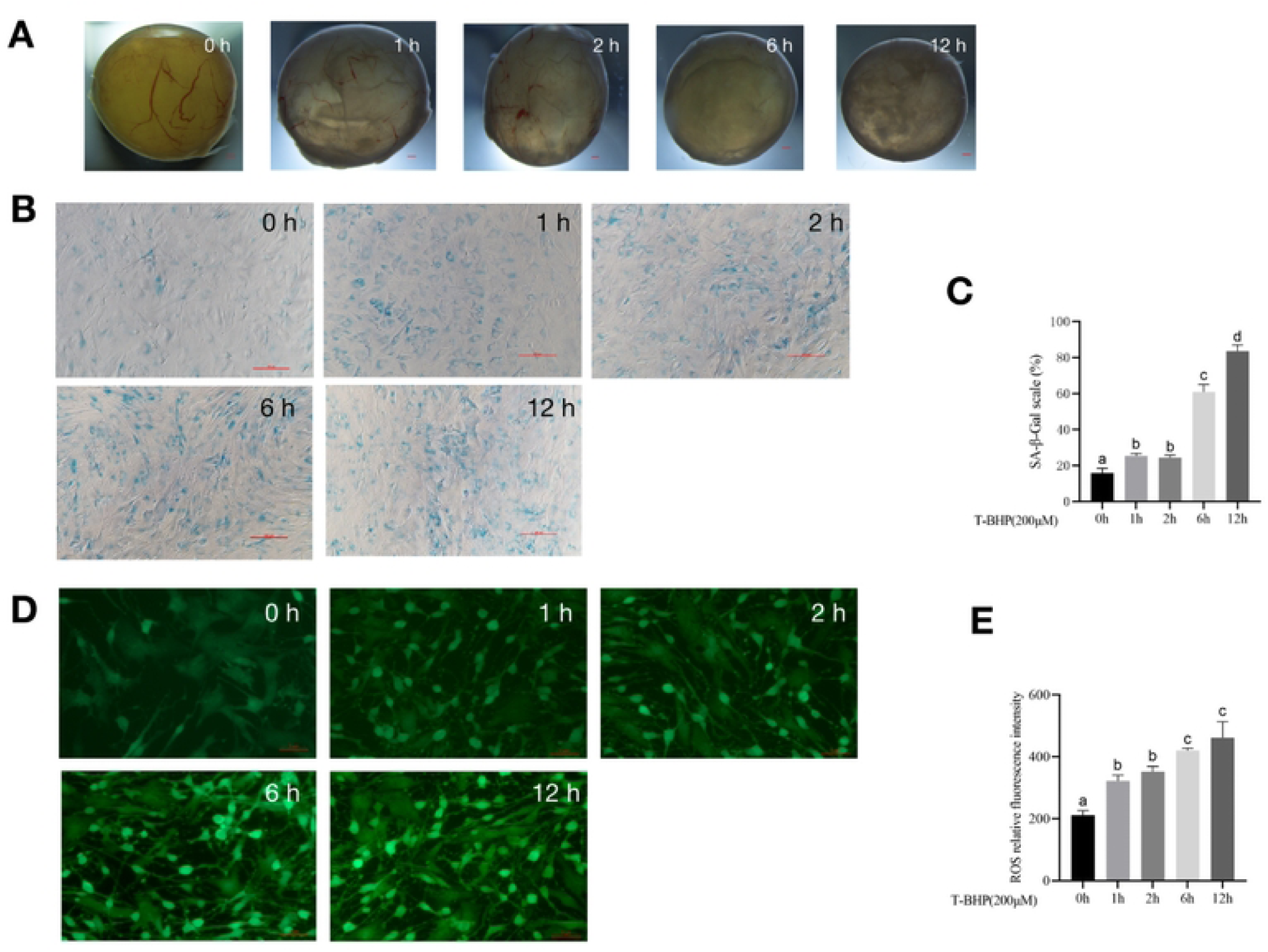
The optimal time of follicular senescence induced by t-BHP. (A) Treating with 200 μM t-BHP, and culturing antral follicles in vitro for 1 h, 2 h, 6 h, and 12 h. Bars represent 10 μm. (B) SA-β-Gal staining of granulosa cells at different time points with 200 μM t-BHP treatment. Bar represents 20 μm. (C) Statistics of positive cells after different time of t-BHP treatment. P< 0.05 (D) ROS fluorescence intensity staining in follicles of each group treated with 200 μM t-BHP at different times. Bar represents 20 μm. (E) Relative ROS levels after different t-BHP treatment times P<0.05.

SA-β-Gal staining of follicular granulosa cells in each group revealed no significant changes in cells in the 1 h and 2 h groups, with a small percentage of cells showing blue staining and a positive cell rate of less than 40%. Compared with the control group, the positive cells in the 6-h group were looser and larger, and the positive staining rate was up to 60% in five randomly selected fields, although cellular senescence occurred in part. However, in the 12-h group, the cells were polygonal, dilated, and flattened, with a decreased refractive index, and most of the cells were senescent, with a positive staining rate of more than 80% in five random fields (Fig 2B, C).

The results of ROS fluorescence staining showed that there was weak ROS staining in the control group, and staining was greater in the 1-h and 2-h group, but there was no significant difference between the two groups. The expression levels of ROS in the 1-h and 2-h groups were higher than those in the control group, and the ROS level increased significantly after 6 h and 12 h. Although the differences between groups were not significant, they all increased significantly over time, with strong fluorescence staining of granulosa cells and poorer cell status in the 12-h group (Fig 2D, E). Under the experimental conditions, the optimal time to induce follicular senescence was 6 h with 200 μM t-BHP.

### The progesterone-to-estrogen ratio increased with time after t-BHP-induced follicular senescence

The progesterone-to-estrogen ratio increased with time after t-BHP-induced follicular senescence. Progesterone can be synthesized in granulosa and membrane cells of follicles, and a low concentration of progesterone contributes to the formation of the LH peak, thus inducing ovulation in mature follicles. Estrogen stimulates granulosa cell proliferation and prevents apoptosis. The results of hormone level determination showed that the concentration of E_2_ decreased with an increase in treatment time, and the change in P_4_ concentration was the opposite; the ratio of P_4_/E_2_ was higher than that of the control group (Table 4).

**Table 4.** The changes of hormone level in follicles.

| Processing time | Progesterone concentration<br>(pmol/L) | Estradiol<br>concentration<br>(pmol/L) | $P_4/E_2$ |
| --- | --- | --- | --- |
| 0 h | 1319.8718±35.13<sup>a</sup> | 93.9583±14.59<sup>a</sup> | 14.69±2.10<sup>a</sup> |
| 1 h | 1316.0256±84.39<sup>a</sup> | 65.0232±7.87<sup>b</sup> | 20.4973±1.08<sup>a</sup> |
| 2 h | 1177.5641±1.28<sup>a</sup> | 62.2454±9.03<sup>b</sup> | 19.7495±2.90<sup>a</sup> |
| 6 h | 1319.8718±39.74<sup>a</sup> | 67.4537±3.61<sup>a</sup> | 19.6361±0.77<sup>a</sup> |
| 12 h | 1201.9231±21.41<sup>a</sup> | 62.5926±4.25<sup>b</sup> | 19.4177±1.59<sup>a</sup> |

### Effect of 6-h treatment with t-BHP on the expression of genes and proteins in porcine follicles

The gene expression levels of *Foxo1, P53, Caspase-3*, and *SOD* were determined by qRT-PCR. Normalized to GAPDH, the gene expression of *Foxo1, P53*, and *Caspase-3* was markedly increased by t-BHP treatment. However, t-BHP partially attenuated the expressions of *SOD* (Fig 3A, B, C, D).

**Fig 3.**
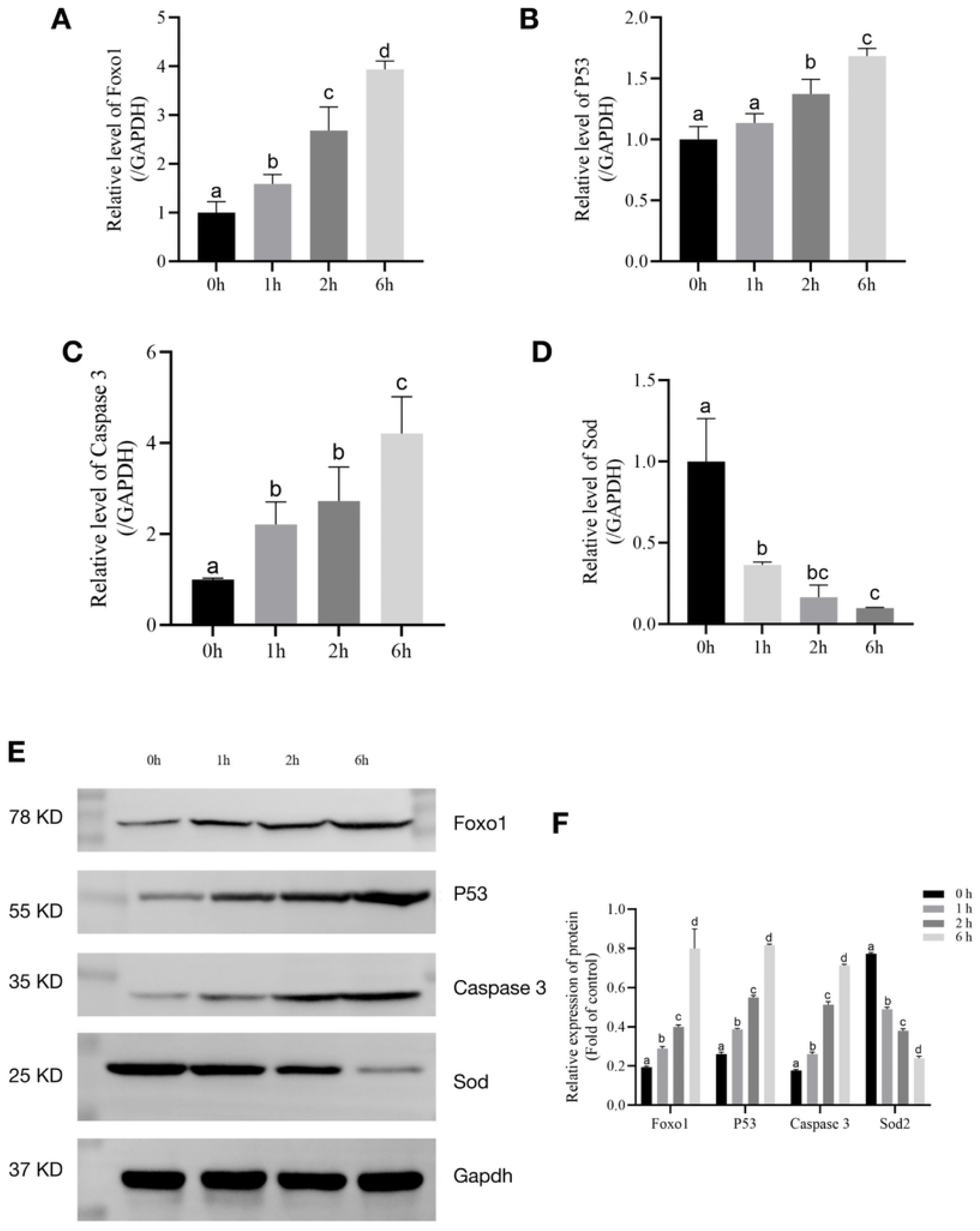
The relative expression levels of *Foxo1, P53, Caspase-3*, and *SOD* after t-BHP treatment for 6 h. (A) Relative expression of mRNA level of *Foxo1* in t-BHP-induced follicles. (P< 0.05). (B) Relative expression of mRNA level of *P53* in t-BHP-induced follicles. (P< 0.05). (C) Relative expression of mRNA level of *Caspase-3* in t-BHP-induced follicles. (P< 0.05). (D) Relative expression of *SOD* in t-BHP-induced follicles. (P< 0.05). (E) Relative protein expression levels of *Foxo1, P53, Caspase-3, SOD*2, and GAPDH in follicles. (F) Statistical analysis of *Foxo1, P53, Caspase-3*, and *SOD*2 in follicles. (P< 0.05)

The protein expression of *Foxo1, P53, Caspase-3*, and *SOD* was determined by western blot analysis. As shown in Fig 3E and F, t-BHP treatment elevated the expression levels of *P53, Caspase-3*, and *Foxo1* compared with the control group. We also found that t-BHP significantly decreased *SOD* protein levels. Taken together, these data implied that t-BHP (200 μM) treatment for 6 h could effectively accelerate senescence through *Foxo1, P53, Caspase-3*, and *SOD2*.

## Discussion

Ovarian failure can lead to infertility, climacterism, or synd. At present, female reproductive aging animal models have gene mutation animal models, such as FSHR knock-out animal models[22]; and chemical damage animal models, such as D-galactose-induced female reproductive senescence model[23]. Although these animal models have shown successful induction of follicular failure and impaired reproductive cycle, knowledge on their effect on sows is limited, and samples from high-parity sows are difficult to obtain; therefore, it is very difficult to carry out thorough studies in pigs by gene knockout and in vivo injection. While other animal models, such as cellular senescence[18], can also lead to multiple secretory phenotypes of senescence, it promotes inflammatory reactions in the body different from the aging of the ovary; therefore, it cannot simulate the real physiological state of aging of the ovary.

Ovarian aging is the basis of reproductive aging. Female mammals are born with a limited number of primordial follicles. Oxidative stress is a prerequisite for luteinization[24] and disrupts the microenvironment within the follicles[25]. Oxidative stress plays an important role in reproductive ovulation, during which oxidative damage gradually accumulates in related cells. Therefore, this study established a simple and easy-to-operate aging model, that is, t-BHP-stimulated follicles, to establish an in vitro aging model.

Ovarian function mainly manifests as endocrine function and follicular atresia[26]. E_2_ and P_4_ levels and granulosa cell senescence are classical criteria for ovarian senescence[27]. This study demonstrated that in vitro culturing of intact antral follicles within 36 h caused atresia, although morphology remained intact; Assessment of cell status revealed that cell viability was higher within 24 h and E_2_ and P_4_ hormone levels were also at normal levels, thus establishing that oxidative stress stimulation of follicles within 24 h of in vitro culture induces senescence.

The accumulation of oxidative stress has long-term effects on follicular failure[28, 29]. Follicles were cultured in vitro with t-BHP at a final concentration of 200 μM. The results showed that ROS levels in the treated group were significantly higher than those in the control group, showing a time-dependent increasing trend and impaired granulosa cell function resulting in follicular atresia. Additionally, follicular granulosa cell morphology significantly changed after 6 h of treatment, with SA-β-Gal staining showing that positive granulosa cells were mostly stained after 6 h. Detection of related hormones showed high progesterone and estrogen levels, indicating that 200 μM t-BHP successfully stimulated follicles for 6 h to accelerate the ovarian aging phenotype.

The intracellular antioxidant system is involved in various oxidation–reduction reactions in organelles and is activated by a low concentration of ROS. Elevated oxidative stress in cells is the leading cause of cellular [30, 31]and ovarian aging[32]. *SOD* is one of the most important antioxidant enzymes in the body, and its expression levels increase following oxidative stress[33]. Based on the findings of the present study, oxidative stress significantly increased the relative expression levels of *P53, Foxo1*, and *Caspase-3* genes and proteins in follicles. The *P53* pathway is an important signaling pathway involved in aging[34]. The expression of *P53* and *SOD*2 proteins is low in young animal tissues, and its expression increases with age. The *Caspase-3* [35]pathway is considered important for activating apoptosis, and the regulation of gene transcription is associated with the initiation of senescence. *Foxo1* induces cellular stress resistance. In our study, western blotting showed that t-BHP stimulation increased *P53, Caspase-3*, and *Foxo1* levels and decreased *SOD* levels. Some evidence supports this result: the *Foxo1* pathway may be an important regulator of cellular senescence.

T-BHP effectively induced oxidative stress, dysregulation of hormone levels, and follicular damage, causing changes in the *SOD, P53, Caspase-3*, and *Foxo1* genes and proteins associated with them. Based on these results, the continuous treatment of follicles with 200 μM t-BHP for 6 h within 24 h of in vitro culture may be an effective model for the establishment of ovarian senescence. In this study, we established a model of oxidative stress in T-BHP-cultured porcine follicles to simulate ovarian senescence. This lays the foundation for subsequent studies on the decline in reproductive ability caused by reproductive senescence in pigs.

## Conclusion

T-BHP effectively induced oxidative stress, dysregulation of hormone levels, and follicular damage, causing changes in the *SOD, P53, Caspase3*, and *Foxo1* genes and proteins associated with them. Based on these results, the continuous treatment of follicles with 200 μM t-BHP for 6 h within 24 h of in vitro culture may be an effective model for the establishment of ovarian senescence.

## Acknowledgments

We would like to thank Dr. Junjie Li for helping with proofreading the manuscript.

## Author Contributions

**Conceptualization:** Peihua Shi.

**Formal analysis:** Peihua Shi, Chenyu Tao.

**Funding acquisition:** Chenyu Tao.

**Investigation:** Peihua Shi.

**Methodology:** Peihua Shi, Shunran Zhao.

**Project administration:** Chenyu Tao.

**Supervision:** Wei Xia, Junjie Li.

**Visualization:** Peihua Shi, Jinchun Gao.

**Writing – original draft:** Peihua Shi, Chenyu Tao.

## Notes

### Competing Interest Statement

The authors have declared no competing interest.

